# Prime Editing Enables High-Efficiency Correction of the Ryr1 T4706M Mutation: A Promising Therapeutic Approach for RyR1-Related Myopathies

**DOI:** 10.1101/2025.09.12.675683

**Authors:** Kelly Godbout, Joël Rousseau, Geoffrey Canet, Jacques P. Tremblay

## Abstract

Prime editing has emerged as a powerful genome-editing tool for precise correction of pathogenic mutations, offering a promising therapeutic approach for genetic myopathies. Here, we evaluate the correction efficiency of the T4706M mutation in the *Ryr1* gene, which is implicated in severe skeletal muscle dysfunction. Using an optimized epegRNA design and RNA electroporation, we achieved a remarkable 80% editing efficiency in immortalized C2C12 myoblasts and 37% correction in primary myoblasts derived from the *RYR1*^*TM/TM*^ mouse model. Our results demonstrate that the PE6 prime editing strategy, combined with rationally designed epegRNAs, significantly enhances editing efficiency in unselected cell populations. These findings establish a critical ex vivo foundation for the development of in vivo Ryr1 gene therapy in preclinical mouse models. They also provide a validated editing design that can support delivery-focused applications in both academic and industry settings.

## Introduction

Prime editing holds significant therapeutic potential for rare genetic diseases by precisely correcting pathogenic mutations at the genomic level. This gene therapy technology employs a SpCas9 nickase (H840A) fused to an engineered reverse transcriptase (RT), whose complex is directed to a precise genomic site by a prime editing guide RNA (pegRNA) [1]. The pegRNA comprises a spacer sequence for genomic targeting, a scaffold for Cas9 binding, a primer binding site (PBS) to initiate reverse transcription, and a reverse transcriptase template (RTT) that encodes the intended sequence modifications [2,3]. To enhance molecular stability, engineered pegRNAs (epegRNAs) incorporate structured elements that mitigate exonuclease-mediated degradation [4]. Several prime editing strategies have been developed [5-7]. In this study, we used PE3, which includes an additional nicking sgRNA (nsgRNA), and PE6, which is a more compact form of the prime editor using a different RT [8]. It is worth noting that, in our study, an additional nsgRNA was included in this strategy. However, we simply refer to it as PE6 throughout the manuscript for ease of reference.

Given its capacity for precise genetic correction, prime editing emerges as a relevant approach for correcting mutations in the *RYR1* gene, which is implicated in several congenital myopathies [9,10], including malignant hyperthermia [11,12], central core disease [13], multi-mini core disease [14], centronuclear myopathy [15], congenital fiber-type disproportion [16] and exertional rhabdomyolysis [17]. The *RYR1* gene encodes ryanodine receptor 1 (RyR1), a key calcium-release channel in skeletal muscle [10]. Pathogenic variants in *RYR1* often lead to impaired calcium homeostasis, ultimately resulting in progressive muscle dysfunction and severe myopathic phenotypes. As many *RYR1* mutations are single-nucleotide variants [18], prime editing represents a precise and targeted therapeutic approach for their correction [19]. In addition to repairing pathogenic mutations, this genome-editing technology has been employed to introduce a protective mutation in the *RYR1* gene, thereby restoring the function of the leaky channel caused by disease-associated variants [20].

While prime editing of RYR1 mutations has been reported by our group in human cells [19], translation to preclinical in vivo models requires prior optimization in the relevant species and cell types. The murine Ryr1 locus differs from its human counterpart, and pegRNA design must be adapted accordingly. Even small sequence variations can substantially impact editing efficiency. Moreover, primary myoblasts are notoriously difficult to transfect and edit, making ex vivo correction a critical step before gene delivery methods can be developed and validated in vivo.

In this study, we assessed the feasibility of correcting the pathogenic T4706M (TM) mutation in the murine Ryr1 gene using prime editing. This variant is the murine equivalent of the human T4709M mutation, which is associated with severe skeletal muscle dysfunction, including proximal muscle weakness, ophthalmoplegia, scoliosis, and respiratory impairment [21]. It was selected for its clinical relevance and the availability of a well-characterized mouse model (*Ryr1*^*TM/TM*^) that recapitulates key pathophysiological features of the disease [22]. We achieved 37% correction in unselected primary myoblasts derived from this model, and an unprecedented 80% editing efficiency in immortalized C2C12 cells. These results were obtained by electroporation of optimized PE components and rational pegRNA design. Together, these findings establish a robust ex vivo editing strategy that lays the groundwork for future in vivo correction of Ryr1 in preclinical models of RyR1-related myopathies.

## Material and Methods

### 4.1 Plasmids

The plasmids pCMV-PE2 (prime editor), pU6-tevopreq1-GG-acceptor (used for epegRNA construction), and pCMV-PE6a were obtained from AddGene (Cambridge, MA, USA) under accession numbers #132775, #174038, and #207851, respectively. Cloning into the pU6-tevopreq1-GG-acceptor plasmid was performed following the protocol established by Anzalone et al. [1] to generate epegRNA-expressing constructs. Plasmid #174038 was further modified by our team to introduce a second U6 promoter along with a cloning site for insertion of a nicking sgRNA, enabling compatibility with the PE3 strategy. This resulting construct is referred to as the epegRNA-nsgRNA plasmid (Supplementary Files). Oligonucleotides used for the design and cloning of epegRNAs and nsgRNAs were synthesized by IDT Inc. (Coralville, IA, USA).

In our prior work, we explored various design parameters including different PBS and RTT lengths, as well as PAM disruption and the addition of silent mutations to reduce re-editing, and identified an optimized configuration for the human *RYR1* T4709M mutation [19]. We then adapted this design to the mouse genome for the homologous mutation (*Ryr1* T4706M). The spacer, PBS, RTT, and nsgRNA sequences used in this study are listed in Table S1.

### 4.2 Cell Culture

C2C12 cells (passage 4) were cultured in DMEM medium (Wisent Inc., Saint-Jean-Baptiste, QC, Canada) supplemented with 10% FBS (Wisent Inc.), and 1% penicillin–streptomycin (Wisent Inc.). Cells were kept at 37 °C with 5% CO_2_ in a humidified incubator.

Primary myoblasts from the *Ryr1*^*TM/TM*^ mouse model muscle were cultured in DMEM medium (Wisent Inc.) supplemented with 20% FBS (Wisent Inc.), and 1% penicillin–streptomycin (Wisent Inc.). Cells were kept at 37 °C with 5% CO_2_ in a humidified incubator.

### 4.3 Primary Myoblast Culture from Adult Mice

#### Dissection and Tissue Collection

A 14-month-old female *Ryr1*^*TM/TM*^ mouse on a C57BL/6 background was used in this study. The animal was euthanized by CO_2_ inhalation and subsequently immersed in 70% ethanol for at least 10 minutes. The excess ethanol was blotted using sterile gauze before making a dorsal skin incision with sterile scissors. The skin was peeled back to expose the hindlimbs, ensuring minimal contamination with fur. The mice were placed on a clean sterile gauze, and excess adipose tissue was removed using fine forceps. Muscle tissue was carefully excised from the paraspinal region and hindlimbs using fine scissors, avoiding connective and adipose tissue contamination. Excised muscles were immediately transferred to a 50 mL Falcon tube containing 20 mL of ice-cold sterile Hank’s Balanced Salt Solution (HBSS).

#### Enzymatic Digestion

Collected muscle tissues were minced into small fragments using sterile scissors in a 10 cm Petri dish. Enzymatic digestion was performed using collagenase (2 mg/mL) and dispase (1 mg/mL) dissolved in 20 mL of HBSS. The solution was filtered through a 0.2 μm sterile filter into a sterile trypsinization unit. Minced muscle fragments were transferred into the unit using a cut-tip pipette, and the digestion mixture was incubated at 37°C for 1 hour with agitation.

#### Cell Isolation and Culture

Following digestion, enzymatic activity was neutralized by adding 20 mL of DMEM (Wisent Inc.) supplemented with 10% FBS (Wisent Inc.) and 1% penicillin-streptomycin (Wisent Inc.). The suspension was transferred to a 50 mL Falcon tube and centrifuged for 1 min at 100 rpm to remove large debris. The supernatant was collected and centrifuged at 680 rpm for 3 min, followed by careful aspiration of the supernatant. The pellet was washed with 40 mL of HBSS and centrifuged at 1400 rpm for 5 min. This washing step was repeated once. The final pellet was resuspended in 30 mL of HBSS and subjected to a low-speed centrifugation step (250 rpm, 2 min) to remove remaining debris. The supernatant was collected and centrifuged at 1400 rpm for 5 min, and the resulting cell pellet was resuspended in 1 mL of DMEM.

#### Coating and Initial Culture

Ten cm Petri dishes were coated with 2 mL of 0.1% sterile gelatin for 1 min, followed by removal of excess gelatin and air-drying for 5 min. The plates were then supplemented with 9 mL of complete DMEM before seeding with 500 μL of the myoblast suspension. Cells were incubated overnight at 37°C with 5% CO_2_.

#### First Passage

After 24 hours, non-adherent cells were collected and centrifuged at 1400 rpm for 5 min. The supernatant was discarded, and the cell pellet was resuspended in 1 mL of DMEM. The suspension was gently pipetted to ensure single-cell dispersion, adjusted to 5 mL with DMEM, and replated on freshly gelatin-coated dishes for further proliferation.

### 4.4 Plasmid Electroporation

Electroporation was carried out using the Neon™ Transfection System (10 μL format, Invitrogen™) along with its corresponding commercial kit, according to the manufacturer’s protocol. For each condition, 100,000 myoblasts were transfected with 1 μg of the pCMV-PE2 plasmid (encoding the prime editor) and 1 μg of the epegRNA-nsgRNA plasmid. Electroporation settings were 1100 V, 20 ms, and two pulses. After transfection, cells were cultured in a 24-well plate with 1 mL of medium per well, which was refreshed 24 hours later. As a control, cells were also electroporated with an eGFP-expressing plasmid.

### 4.5 PE mRNA In Vitro Transcription

The prime editor plasmids (pCMV-PE2 or pCMV-PE6a) were first amplified by PCR to incorporate optimized 5′ untranslated regions (UTRs) and a polyadenylation signal at the 3′ end (see Table S1). The resulting PCR products were purified using the EZ-10 Spin Column PCR Products Purification Kit (Bio Basic, Toronto, ON, Canada) and used as templates for in vitro transcription (IVT). Transcription was carried out with the HiScribe T7 mRNA Kit supplemented with CleanCap Reagent AG (New England BioLabs Inc., Ipswich, MA, USA), replacing UTP entirely with N1-Methylpseudouridine-triphosphate (TriLink Biotechnologies Inc., San Diego, CA, USA). The reaction was incubated at 37 °C for 3 hours and 30 minutes. Afterwards, the volume was adjusted to 50 μL with nuclease-free water, and 2 μL of DNase I was added to degrade the template DNA, followed by an additional 15-minute incubation at 37 °C. The synthesized mRNAs were then purified using the Monarch RNA Clean-up Kit (500 μg format, New England BioLabs Inc.), eluted in 1 mM sodium citrate (pH 6.4), and quantified using a BioDrop spectrophotometer (Cambridge, UK). The resulting PE mRNAs were stored at −80 °C until use.

### 4.6 RNA Electroporation

A total of 1 μg of PE or PE6 mRNA, 4.6 μg of epegRNA (chemically synthesized by IDT Inc., Coralville, IA, USA), and 1.8 μg of nsgRNA (also synthesized by IDT Inc.) were delivered into 100,000 human myoblasts via electroporation using the Neon Transfection System (10 μL tips; settings: 1100 V, 20 ms, 2 pulses). Following electroporation, cells were transferred to a 24-well plate containing 500 μL of homemade culture medium per well. Control conditions included electroporation with eGFP mRNA produced by IVT. The culture medium was replaced 24 hours post-electroporation with 1 mL of fresh medium. Genomic DNA was extracted from the cells 48 hours after transfection.

### 4.7. Genomic DNA Preparation and PCR Amplification

Myoblasts were washed in their wells using 500 μL of PBS. Subsequently, 100 μL of Di-rectPCR Lysis Reagent (Viagen Biotech Inc., Los Angeles, CA, USA) mixed with 1 μL of proteinase K (20 mg/mL) was added to each well. Cell lysates were incubated for 2 hours at 56 °C, followed by heat inactivation for 45 minutes at 85 °C. Samples were then centrifuged at 13,500 rpm for 5 minutes. From the supernatant (genomic DNA), 1 to 2 μL was used as template for PCR amplification. The thermal cycling conditions were: initial denaturation at 98 °C for 30 seconds; 35 cycles of 98 °C for 10 seconds, 58 °C for 20 seconds, and 72 °C for 30 seconds; followed by a final extension step at 72 °C for 5 minutes. All PCR reactions were performed using Phusion™ High-Fidelity DNA Polymerase (ThermoFisher Scientific Inc., Waltham, MA, USA). Primer sequences are provided in Table S1.

### 4.8 Sanger Sequencing

Sanger sequencing of PCR amplicons was performed at the sequencing platform of the CHU de Québec Research Center (https://sequences.ulaval.ca/murin/servseq.pageaccueil). Polymerization was carried out using an internal primer (Table S1) and the BigDye™ Terminator v3.1 (ThermoFisher Scientific Inc. (Waltham, MA, USA The resulting sequences were analyzed for editing efficiency using the online tool EditR (https://moriaritylab.shinyapps.io/editr_v10/), accessed between 1 September 2022 and 20 December 2024 [23].

#### 4.9 Statistical Analysis

Data were analyzed using the GraphPad PRISM 10.3.0 software package (Graph Pad Software Inc., La Jolla, CA, USA).

## Results

An epegRNA adapted for the mouse genome was designed and constructed based on the one used in our previous study on human *RYR1*^*TM/TM*^ myoblasts [19]. A silent mutation adjacent to the intended correction was incorporated to enhance editing efficiency and facilitate tracking of the editing rate in wild-type cells.

The efficiency of the epegRNA was first evaluated in C2C12 myoblasts using plasmid DNA delivery, resulting in 11% editing of the adjacent silent mutation with the PE3 strategy. When electroporating RNA instead of plasmid DNA, the editing rate increased to 54% with the PE3 strategy. Furthermore, employing PE6, a more compact version of prime editing, led to an editing rate of 70% (Figure 2a). Notably, these cells were neither sorted nor selected, meaning that the reported editing percentage represents the raw efficiency observed in the entire cell population. Different ratios of the prime editing components were also tested (Figure 2b). Doubling the quantity of all the components resulted in a slightly higher editing rate, reaching 80% editing, while not being significantly different.

**Figure 1.**
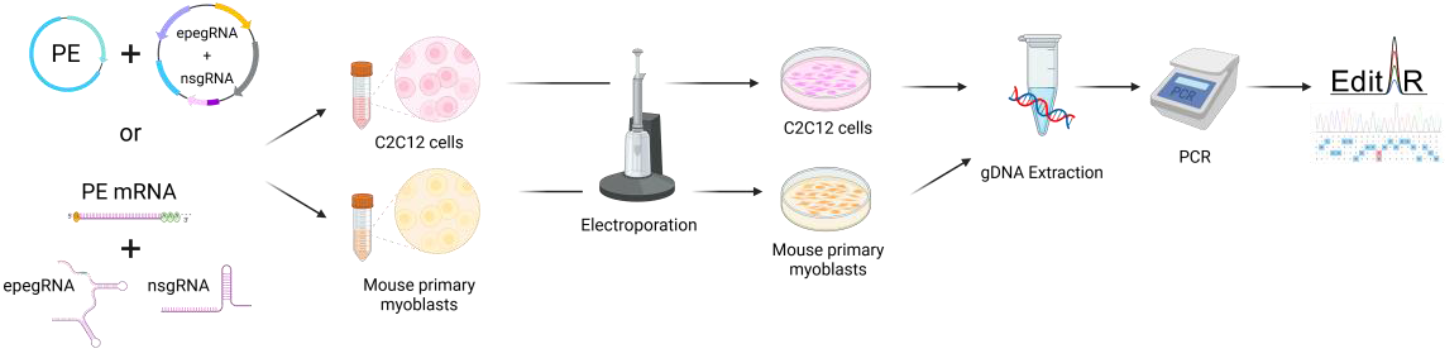
Schematic representation of the prime editing workflow used to correct the *Ryr1* T4706M mutation in C2C12 cells and primary mouse myoblasts. Prime editing components (plasmids or RNAs) were introduced by electroporation using the Neon Transfection System. Genomic DNA was extracted, amplified by PCR, and analyzed by Sanger sequencing using the EditR tool to quantify editing efficiency.

**Figure 2.**
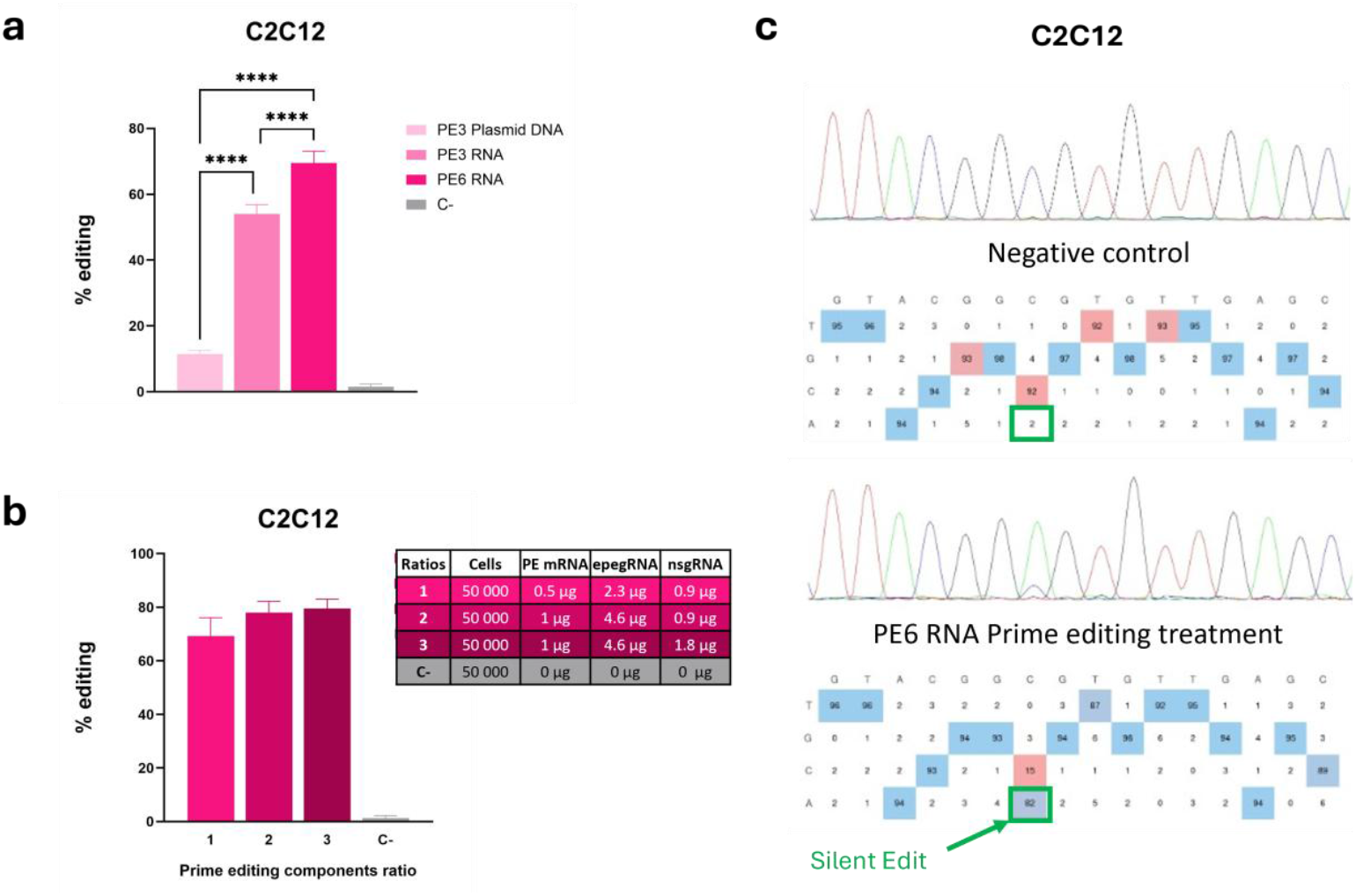
(a) Insertion in wild-type C2C12 myoblasts of a silent mutation adjacent to the eventual pathogenic T4706M mutation in the codon 4706 of the *Ryr1* gene by Prime editing. N = 7 replicates were performed for the PE3 plasmid DNA and negative control (C-) groups. The experiment for the PE3 RNA and PE6 RNA group was performed in biological duplicates. One-way ANOVA was used as a statistical test and Tukey’s multiple comparison test was performed. ^****^ represents a p-value < 0.0001. All conditions are significantly different from the negative control (C-). (b) Insertion in wild-type C2C12 myoblasts of a silent mutation adjacent to the eventual pathogenic T4706M mutation in the codon 4706 of the *Ryr1* gene by Prime editing using different ratios of the prime editing components. N = 8 replicates were performed for the Ratio 1 group. The experiment for the Ratio 2 and Ratio 3 groups was performed in biological duplicates, while C-was performed with N=4 replicates. One-way ANOVA was used as a statistical test and Tukey’s multiple comparison test was performed. Ratio 1, 2 and 3 did not led to a significant difference in editing efficiency. All conditions are significantly different from the negative control (C-). (c) Example of sequencing results. Sequences were analyzed with the EditR online program (https://moriaritylab.shinyapps.io/editr_v10/) [23] to analyze editing efficiencies. In the top of the panel is a representative visualization of the unedited sequence of C2C12 myoblasts. In the bottom of the panel is a representative result of editing efficiencies by PE6 delivered by RNA for the insertion in wild-type C2C12 myoblasts of a silent mutation (green) adjacent to the eventual pathogenic T4706M mutation in the codon 4706 of the *Ryr1* gene.

To assess the correction efficiency of the TM mutation, primary myoblast cultures were established from the muscles of the *RyR1*^*TM/TM*^ mouse model. Electroporation of plasmid DNA using the PE3 strategy resulted in a modest 4% correction rate. However, when the PE6 strategy was delivered via RNA electroporation, the TM mutation was corrected in 37% of the cell population (Figure 3b).

**Figure 3.**
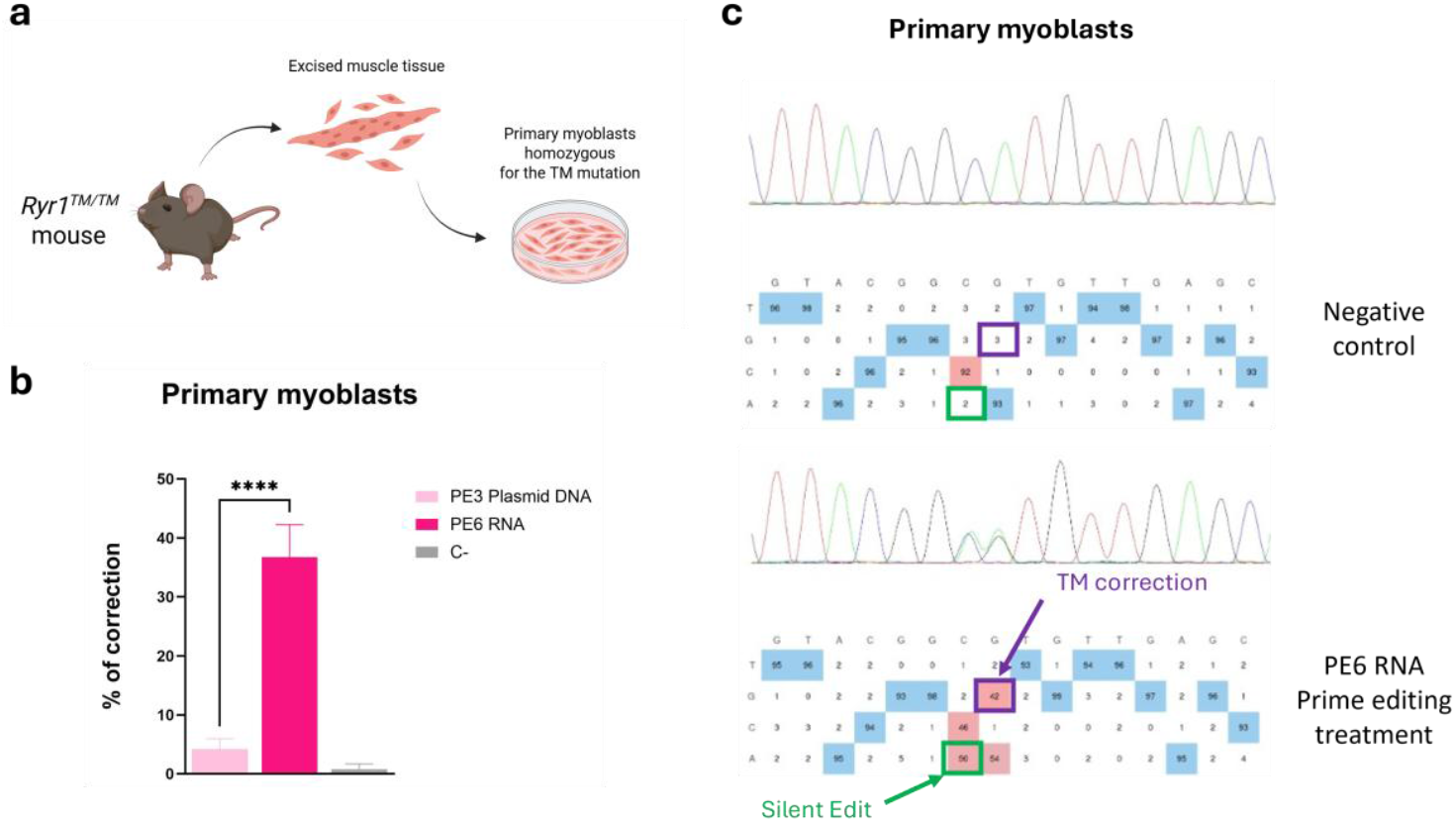
(a) Primary myoblasts homozygous for the T4706M (TM) mutation were obtained from excised muscle tissue from a *Ryr1*^*TM/TM*^ mouse. (b) Correction of the T4706M (TM) mutation in the *Ryr1* gene in primary myoblasts cultured from the muscles of the *Ryr1*^*TM/TM*^ mouse model. PE3 plasmid DNA and PE6 RNA delivery were assessed. N = 4 replicates were performed. One-way ANOVA was used as a statistical test and Tukey’s multiple comparison test was performed. ^***^ represents a p-value < 0.0001. The PE6 RNA group was also significantly different from the negative control (C-), but not the PE3 plasmid DNA group. (c) Example of sequencing results. Sequences were analyzed with the EditR online program (https://moriaritylab.shinyapps.io/editr_v10/) [23] to analyze editing efficiencies. In the top of the panel is a representative visualization of the unedited sequence of primary myoblasts cultured from the muscles of the *Ryr1*^*TM/TM*^ mouse model. In the bottom of the panel is a representative result of editing efficiencies by PE6 delivered by RNA for the correction of the T4706M (TM) mutation in the *Ryr1* gene in primary myoblasts cultured from the muscles of the *Ryr1*^*TM/TM*^ mouse model (purple), and also for the insertion of a silent mutation (green) adjacent to the pathogenic T4706M mutation in the codon 4706 of the *Ryr1* gene.

## Discussion

Our findings demonstrate that our prime editing design can efficiently correct the T4706M mutation in the *Ryr1* gene with a correction rate of 37% in primary myoblasts when using RNA delivery and the PE6 strategy. The editing efficiency observed in immortalized C2C12 cells further supports the robustness of this approach, particularly with the transition from PE3 to PE6 and from DNA to RNA electroporation, which significantly enhanced the correction rate. These results highlight the potential of prime editing for precise genomic modifications in myoblasts.

Our results are particularly noteworthy due to the high editing efficiency achieved in myoblasts without requiring multiple treatments or selection-based enrichment through antibiotic resistance or fluorescence-based sorting. Using a single treatment, our approach yielded up to 80% editing in immortalized C2C12 cells and 37% in primary myoblasts in unselected populations, highlighting the intrinsic efficiency of our optimized epegRNA design and RNA delivery method. Notably, our editing efficiency exceeds previously reported outcomes in myoblasts without selection, such as those by Halega et al. [24], who achieved 25% editing at the *HEK3* locus, and Mbakam et al. [25], who reported 22% correction in the *DMD* gene. Halega et al. employed a PE-VLP system (Nanoscribes) to deliver PE3 components [24], while Mbakam et al. [25] used DNA electroporation to introduce PE5 components—while Mbakam et al. used DNA electroporation to introduce PE5 components—an approach similar to PE3 but incorporating co-expression of MLH1dn, which encodes a mismatch repair inhibitory protein. In contrast, our study significantly enhanced prime editing efficiency by utilizing RNA electroporation and transitioning to the PE6 system. This underscores the robustness of our strategy, making it particularly relevant for therapeutic applications where selection-based enrichment is not feasible in vivo.

Importantly, our current strategy also outperforms our own previous work on the same mutation in the human genome context. The improved editing efficiencies showed here can be attributed to two key factors: the use of the latest PE6 prime editor, and the better quality and responsiveness of the cellular models employed. In our previous study [19], the TM homozygous human myoblast cell line was generated by four consecutive electroporation treatment, possibly affecting cell health and compromising responsiveness to editing. In contrast, unmodified, commercially available C2C12 cells were used here. These improvements—both in editing machinery and cellular context—enabled us to achieve high correction rates (80% in C2C12 cells) after a single RNA electroporation, without selection.

Freshly isolated primary mouse myoblasts were also used. While notable correction rates were obtained with those cells (37% of correction), the use of primary myoblast cultures derived from adult *Ryr1*^*TM/TM*^ mice instead of neonatal mice is a key limitation of our study. It is well-established that myoblast cultures from adult mice are often of lower quality compared to those from neonatal mice, primarily due to a higher proportion of fibroblasts in the culture [26]. This is particularly relevant to our study since *Ryr1* expression is restricted to myoblasts, meaning that fibroblasts do not contribute to the observed editing efficiency [19]. Consequently, the correction rate of 37% in primary myoblasts may be an underestimation, as a higher proportion of myoblasts in neonatal-derived cultures would likely result in even greater editing efficiency.

While no overt toxicity or abnormal phenotypes was observed following RNA electroporation and prime editing, we acknowledge that off-target or bystander effects remain a potential concern. Future work will be required to comprehensively assess genome-wide specificity in both in vitro and in vivo settings. The use of rationally designed epegRNAs and transient delivery via RNA electroporation may help mitigate such risks.

Given the promising correction rates observed in this study, future research should focus on the in vivo delivery of the prime editing machinery. Efficient and targeted delivery methods, such as lipid nanoparticles (LNPs) [27], adeno-associated viruses (AAVs) [28] or extracellular vesicles [29], hold promise to facilitate muscle-specific gene correction. Although in vivo correction is the next logical step, developing effective delivery systems remains a complex, multidisciplinary challenge. By providing an efficient, species-specific editing strategy in disease-relevant murine cells, our study supports the efforts of teams specializing in vector development and enables translational research to proceed more efficiently. Beyond delivery optimization, further studies will be required to assess functional improvements in muscle physiology and to define the precise editing threshold necessary for therapeutic benefit. These steps will be essential for the ultimate clinical translation of prime editing therapies for RyR1-related myopathies. Overall, this work represents a key preclinical milestone toward achieving functional Ryr1 gene correction in vivo.

## Conclusion

The high editing efficiencies achieved in both C2C12 cells and primary myoblasts highlight the strong potential of prime editing for therapeutic applications. With a correction rate of up to 80% in immortalized myoblasts and 37% in primary myoblasts, our findings suggest that prime editing could reach therapeutically relevant levels for treating RyR1-RM. These results provide a solid foundation for the development of prime editing as a viable gene therapy approach for those rare diseases which currently have no cure.

## Author Contributions

Conceptualization, K.G., J.R. and J.P.T.; methodology, K.G. and J.R.; validation, K.G.; formal analysis, K.G.; investigation, K.G.; writing—original draft preparation, K.G and G.C..; writing—review and editing, K.G., G.C., J.R. and J.P.T.; visualization, K.G.; supervision, J.R. and J.P.T.; funding acquisition, J.P.T. All authors have read and agreed to the published version of the manuscript.

## Institutional Review Board Statement

The study was conducted in accordance with the Decla-ration of Helsinki and approved by the Ethics Committee of the Centre de recherche du CHU de Québec —Université Laval (approval code: 130780).

## Informed Consent Statement

Not applicable.

## Data Availability Statement

The original contributions presented in this study are included in the article/Supplementary Materials. Further inquiries can be directed to the corresponding author.

## Acknowledgments

The authors utilized ChatGPT-4.0 to enhance the readability and clarity of the manuscript. Following its use, the authors thoroughly reviewed and edited the content as necessary, ensuring its accuracy and taking full responsibility for the final version of the publication.

