## Supplementary material for "Prime Editing Enables High-Efficiency Correction of the Ryr1 T4706M Mutation: A Promising Therapeutic Approach for RyR1-Related Myopathies": p2U6-pegRNA_mRyr1_TM-sequence.pdf

### p2U6-pegRNA_mRyr1_TM (2710 bp)

```
tacgcgcagaaaaaaaggatctcaagaagatcctttgatcttttctacggggtctgacgctcagtggaacgaaaactcacgttaagggattttggtcatgagattat
atgcgcgtctttttttcctagagttcttctaggaaactagaaaagatgccccagactgcgagtcaccttgcttttgagtgcaattccctaaaaccagtactctaata
ori
10 20 30 40 50 60 70 80 90 100

caaaaaggatcttcacctagatccttttaaattaaaaatgaagttttaaatcaatctaaagtatatatgagtaaacttggtctgacagttaccaatgcttaatcagt
gtttttcctagaagtggatctaggaaaatttaatttttacttcaaaatttagttagatttcatatatactcatttgaaccagactgtcaatggttacgaattagtca
AmpR
110 120 130 140 150 160 170 180 190 200 210

gaggcacctatctcagcgatctgtctatttcgttcatccatagttgcctgactccccgtcgtgtagataactacgatacgggagggcttaccatctggccccagtgc
ctccgtggatagagtcgctagacagataaagcaagtaggtatcaacggactgaggggcagcacatctattgatgctatgccctcccgaatggtagaccggggtcacg
AmpR
220 230 240 250 260 270 280 290 300 310 320

tgcaatgataccgcgagatccacgctcaccggctccagatttatcagcaataaaccagccagccggaagggccgagcgcagaagtggtcctgcaactttatccgcct
acgttactatggcgctctaggtgcgagtggccgaggtctaaatagtcgttatttggtcggtcggccttcccggctcgcgtcttcaccaggacgttgaaataggcgga
AmpR
330 340 350 360 370 380 390 400 410 420

ccatccagtctattaattgttgccgggaagctagagtaagtagttcgccagttaatagtttgcgcaacgttgttgccattgctacaggcatcgtggtgtcacgctcg
ggtaggtcagataattaacaacggcccttcgatctcattcatcaagcggtcaattatcaaacgcgttgcaacaacggtaacgatgtccgtagcaccacagtgcgagc
AmpR
430 440 450 460 470 480 490 500 510 520 530

tcgtttggtatggcttcattcagctccggttcccaacgatcaaggcgagttacatgatcccccatgttgtgcaaaaaagcggttagctccttcggtcctccgatcgt
agcaaaccataccgaagtaagtcgaggccaagggttgctagttccgctcaatgtactagggggtacaacacgttttttcgccaatcgaggaagccaggaggctagca
AmpR
540 550 560 570 580 590 600 610 620 630 640

tgtcagaagtaagttggccgcagtgttatcactcatggttatggcagcactgcataattctcttactgtcatgccatccgtaagatgcttttctgtgactggtgagt
acagtcttcattcaaccggcgtcacaatagtgagtaccaataccgtcgtgacgtattaagagaatgacagtacggtaggcattctacgaaaagacactgaccactca
AmpR
650 660 670 680 690 700 710 720 730 740

actcaaccaagtcattctgagaatagtgtatgcggcgaccgagttgctcttgcccggcgtcaatacgggataataccgcgccacatagcagaactttaaaagtgctc
tgagttggttcagtaagactcttatcacatacgccgctggctcaacgagaacgggccgcagttatgccctattatggcgcggtgtatcgtcttgaaattttcacgag
Amp-R
AmpR
750 760 770 780 790 800 810 820 830 840 850
```

```
atcattggaaaacgttcttcggggcgaaaactctcaaggatcttaccgctgttgagatccagttcgatgtaacccactcgtgcacccaactgatcttcagcatcttt
tagtaaccttttgcaagaagccccgcttttgagagttcctagaatggcgacaactctaggtcaagctacattgggtgagcacgtgggttgactagaagtcgtagaaa
```

AmpR

860 870 880 890 900 910 920 930 940 950 960

```
tactttcaccagcgtttctgggtgagcaaaaacaggaaggcaaaatgccgcaaaaaagggaataagggcgacacggaaatgttgaatactcatactcttcctttttc
atgaaagtggtcgcaaagacccactcgtttttgtccttccgttttacggcgttttttcccttattcccgctgtgcctttacaacttatgagtatgagaaggaaaaag
```

AmpR | AmpR...ter

970 980 990 1,000 1,010 1,020 1,030 1,040 1,050 1,060 1,070

```
aatattattgaagcatttatcagggttattgtctcatgagcggatacatatttgaatgtatttagaaaaataaacaaataggggttccgcgcacatttccccgaaaa
ttataataacttcgtaaatagtcccaataacagagtactcgcctatgtataaacttacataaatctttttatttgtttatccccaaggcgcgtgtaaaggggctttt
```

Seq_pU6_F

AmpR promoter

1,080 1,090 1,100 1,110 1,120 1,130 1,140 1,150 1,160 1,170

```
gtgccacctgacgtcgctagctgtacaaaaaagcaggctttaaaggaaccaattcagtcgactggatccggtaccaaggtcgggcaggaagagggcctatttcccat
cacggtggactgcagcgatcgacatgttttttcgtccgaaatttccttggttaagtcagctgacctaggccatggttccagcccgtccttctcccggataaagggta
```

U6 promoter

Pr_Pu6 ...loning | hU6-F

1,180 1,190 1,200 1,210 1,220 1,230 1,240 1,250 1,260 1,270 1,280

```
gattccttcatatttgcatatacgatacaaggctgttagagagataattagaattaatttgactgtaaacacaaagatattagtacaaaatacgtgacgtagaaagt
ctaaggaagtataaacgtatatgctatgttccgacaatctctctattaatcttaattaaactgacatttgtgtttctataatcatgttttatgcactgcatctttca
```

U6 promoter

1,290 1,300 1,310 1,320 1,330 1,340 1,350 1,360 1,370 1,380 1,390

```
aataatttcttgggtagtttgcagttttaaaattatgttttaaaatggactatcatatgcttaccgtaacttgaaagtatttcgatttcttggctttatatatcttg
ttattaaagaacccatcaaacgtcaaaattttaatacaaaattttacctgatagtatacgaatggcattgaactttcataaagctaaagaaccgaaatatatagaac
```

U6 promoter

LKO.1 5’

1,400 1,410 1,420 1,430 1,440 1,450 1,460 1,470 1,480 1,490

```
tggaaaggacgaaacaccgAGGTGGGGCTGGTCACTTAGTTTTAgagctagaaatagcaagttaaaataaggctagtccgttatcaacttgaaaaagtggcaccgag
acctttcctgctttgtggcTCCACCCCGACCAGTGAATCAAAATctcgatctttatcgttcaattttattccgatcaggcaatagttgaactttttcaccgtggctc
```

U6...r | Sp scaffold R

Spacer | Sp scaffold

1,500 1,510 1,520 1,530 1,540 1,550 1,560 1,570 1,580 1,590 1,600

```
tcgGTGCGCTCAACACTCCGTAAGTGACCAGCCcgcggttctatctagttacgcgttaaaccaactagaatttttttaagcttgggccgCTCGAGTTTCCCATGATT
agcCACGCGAGTTGTGAGGCATTCACTGGTCGGgcgccaagatagatcaatgcgcaatttggttgatcttaaaaaaattcgaacccggcGAGCTCAAAGGGTACTAA
```

RTT | PBS | Tevo | U6-2

1,610 1,620 1,630 1,640 1,650 1,660 1,670 1,680 1,690 1,700 1,710

```
CCTTCATATTTGCATATACGATACAAGGCTGTTAGAGAGATAATTAGAATTAATTTGACTGTAAACACAAAGATATTAGTACAAAATACGTGACGTAGAAAGTAATA
GGAAGTATAAACGTATATGCTATGTTCCGACAATCTCTCTATTAATCTTAATTAAACTGACATTTGTGTTTCTATAATCATGTTTTATGCACTGCATCTTTCATTAT
```

U6-2

1,720 1,730 1,740 1,750 1,760 1,770 1,780 1,790 1,800 1,810

```
ATTTCTTGGGTAGTTTGCAGTTTTAAAATTATGTTTTAAAATGGACTATCATATGCTTACCGTAACTTGAAAGTATTTCGATTTCTTGGCTTTATATATCTTGTGGA
TAAAGAACCCATCAAACGTCAAAATTTTAATACAAAATTTTACCTGATAGTATACGAATGGCATTGAACTTTCATAAAGCTAAAGAACCGAAATATATAGAACACCT
```

U6-2

1,820 1,830 1,840 1,850 1,860 1,870 1,880 1,890 1,900 1,910 1,920

```
AAGGACGAAAcaccgCTACATTACAGAGCAGCCCGGTTTAAGAGCTATGCTGGAAACAGCATAGCAAGTTTAAATAAGGCTAGTCCGTTATCAACTTGAAAAAGTGG
TTCCTGCTTTgtggcGATGTAATGTCTCGTCGGGCCAAATTCTCGATACGACCTTTGTCGTATCGTTCAAATTTATTCCGATCAGGCAATAGTTGAACTTTTTCACC
```

nsgRNA | Pr R...RNA

1,930 1,940 1,950 1,960 1,970 1,980 1,990 2,000 2,010 2,020 2,030

```
CACCGAGTCGGTGCTTTTTTTGCGGCCGCCTCGAGgtacctctctacatatgacatgtgagcaaaaggccagcaaaaggccaggaaccgtaaaaaggccgcgttgct
GTGGCTCAGCCACGAAAAAAACGCCGGCGGAGCTCcatggagagatgtatactgtacactcgttttccggtcgttttccggtccttggcatttttccggcgcaacga
```

Pr_pU6 ...loning

2,040 2,050 2,060 2,070 2,080 2,090 2,100 2,110 2,120 2,130 2,140

```
ggcgtttttccataggctccgcccccctgacgagcatcacaaaaatcgacgctcaagtcagaggtggcgaaacccgacaggactataaagataccaggcgtttcccc
ccgcaaaaaggtatccgaggcggggggactgctcgtagtgtttttagctgcgagttcagtctccaccgctttgggctgtcctgatatttctatggtccgcaaagggg
```

pBR322ori-F

ori

2,150 2,160 2,170 2,180 2,190 2,200 2,210 2,220 2,230 2,240

```
ctggaagctccctcgtgcgctctcctgttccgaccctgccgcttaccggatacctgtccgcctttctcccttcgggaagcgtggcgctttctcatagctcacgctgt
gaccttcgagggagcacgcgagaggacaaggctgggacggcgaatggcctatggacaggcggaaagagggaagcccttcgcaccgcgaaagagtatcgagtgcgaca
```

ori

2,250 2,260 2,270 2,280 2,290 2,300 2,310 2,320 2,330 2,340 2,350

```
aggtatctcagttcggtgtaggtcgttcgctccaagctgggctgtgtgcacgaaccccccgttcagcccgaccgctgcgccttatccggtaactatcgtcttgagtc
tccatagagtcaagccacatccagcaagcgaggttcgacccgacacacgtgcttggggggcaagtcgggctggcgacgcggaataggccattgatagcagaactcag
```

ori

2,360 2,370 2,380 2,390 2,400 2,410 2,420 2,430 2,440 2,450 2,460

```
caacccggtaagacacgacttatcgccactggcagcagccactggtaacaggattagcagagcgaggtatgtaggcggtgctacagagttcttgaagtggtggccta
gttgggccattctgtgctgaatagcggtgaccgtcgtcggtgaccattgtcctaatcgtctcgctccatacatccgccacgatgtctcaagaacttcaccaccggat
ori
    2,470     2,480     2,490     2,500     2,510     2,520     2,530     2,540     2,550     2,560

actacggctacactagaagaacagtatttggtatctgcgctctgctgaagccagttaccttcggaaaaagagttggtagctcttgatccggcaaacaaaccaccgct
tgatgccgatgtgatcttcttgtcataaaccatagacgcgagacgacttcggtcaatggaagcctttttctcaaccatcgagaactaggccgtttgtttggtggcga
ori
2,570     2,580     2,590     2,600     2,610     2,620     2,630     2,640     2,650     2,660     2,670

ggtagcggtggtttttttgtttgcaagcagcagat
ccatcgccaccaaaaaaacaaacgttcgtcgtcta
ori
   2,680     2,690     2,700   2,710
```
